# Phosphoserine acidic cluster motifs in the cytoplasmic domains of transmembrane proteins bind distinct basic regions on the μ subunits of clathrin adaptor protein complexes

**DOI:** 10.1101/286633

**Authors:** Rajendra Singh, Charlotte Stoneham, Christopher Lim, Xiaofei Jia, Javier Guenaga, Richard Wyatt, Joel O. Wertheim, Yong Xiong, John Guatelli

**Affiliations:** Department of Medicine, University of California San Diego, La Jolla, California, USA; Department of Molecular Biophysics and Biochemistry, Yale University, New Haven, Connecticut, USA; Department of Chemistry and Biochemistry, University of Massachusetts Dartmouth, Dartmouth, Massachusetts; Department of Immunology and Microbiology, The Scripps Research Institute, La Jolla, CA 92037; The VA San Diego Healthcare System, San Diego, California, USA

**Author notes:** Corresponding authors: Rajendra Singh, John Guatelli.

**Keywords:** clathrin, adaptor proteins, medium subunits, acidic cluster, phosphoserine, HIV-1 Vpu, Furin, Serinc3

## Abstract

Protein trafficking in the endosomal system involves the recognition of specific signals within the cytoplasmic domains (CDs) of transmembrane proteins by clathrin adaptors. One such signal is the phosphoserine acidic cluster (PSAC), the prototype of which is in the endoprotease Furin. How PSACs are recognized by clathrin adaptors has been controversial. We reported previously that HIV-1 Vpu, which modulates cellular immunoreceptors, contains a PSAC that binds to the µ subunits of clathrin adaptor protein (AP) complexes. Here, we show that the CD of Furin binds the µ subunits of AP-1 and AP-2 in a phosphorylation-dependent manner. Moreover, we identify a PSAC in a cytoplasmic loop of the cellular transmembrane Serinc3, an inhibitor of the infectivity of retroviruses. The two serines within the PSAC of Serinc3 are phosphorylated by casein kinase II and mediate interaction with the µ subunits in vitro. The sites of these serines vary among mammals in a manner consistent with host-pathogen conflict, yet the Serinc3-PSAC seems dispensible for anti-HIV activity and for counteraction by HIV-1 Nef. The CDs of Vpu, Furin, and the PSAC-containing loop of Serinc3 each bind the μ subunit of AP-2 (µ2) with similar affinities, but they appear to utilize different basic regions on µ2. The Serinc3 loop requires a region previously reported to bind the acidic plasma membrane lipid phosphatidylinositol-4,5-bisphosphate. These data suggest that the PSACs within different proteins recognize different basic regions on the µ surface, providing the potential to inhibit the activity of viral proteins without necessarily affecting cellular protein trafficking.

## Introduction

Protein sorting within the endosomal system relies in large part on the recognition of short, linear peptide sequences in the cytoplasmic domains of transmembrane proteins (sorting signals) by the coat proteins of transport vesicles [reviewed in (1)]. Heterotetrameric clathrin adaptor protein complexes (the AP complexes) play a prominent role in this process [reviewed in (2)]. The AP complexes bind both clathrin and various sorting signals [reviewed in (3)]. The sorting signals are of several types and are commonly tyrosine-based (conforming to the sequence Yxxϕ, where ϕ denotes a bulky hydrophobic residue), leucine-based (conforming to the sequence ExxxLϕ), or contain phosphoserines flanked by acidic residues (hereafter termed phosphoserine acidic clusters or PSACs)(1-4).

While the structural basis of how tyrosine- and leucine-based sorting signals bind the AP complexes has been defined(5,6), how PSAC signals bind is unclear. Certain studies suggest that PSAC sequences, the prototype of which - SDSEEDE- is found in the cytoplasmic domain of the endoprotease Furin(4), bind to AP complexes via an intermediary adaptor named PACS-1(7). Others have suggested that these acidic sequences bind directly to the medium-sized (µ) subunit of the AP complexes(8,9), a compelling notion given the highly basic nature of the surfaces of the µ subunits(5). In a series of studies culminating in an X-ray cystallographic model, we reported that HIV-1 Nef, a peripheral membrane protein that provides immune evasion by preventing peptide-loaded class I MHC from reaching the cell surface, formed a co-operative ternary complex with the cytoplasmic domain (CD) of the class I MHC α chain and the µ subunit of AP-1 (µ1), a clathrin adaptor involved in transport within the endosomal system(10-12). The interaction of acidic residues both in Nef and in the MHC-I CD with basic regions on µ1 were critical to the formation of this complex: one region (herein basic region 1) participated in a three-way hydrogen-bonding network with residues in both Nef and the MHC-I CD, whereas the other (herein basic region 2) appeared to provide an electrostatic interaction with an acidic cluster (four consecutive glutamic acid residues) on Nef(12). Notably, the acidic cluster on Nef contains no serines.

Recently, we reported that HIV-1 Vpu, a small transmembrane protein that provides immune evasion by modulating several cellular receptors including CD4, BST2, NTB-A, and CCR7(13-16), contains a PSAC of the sequence EDSGNESE in its cytoplasmic domain(17). The serines, when phosphorylated, constitute a phosphodegron that interacts with a β-TrCP/cullin-1 E3 ubiquitin ligase complex(18), but they are also required for a phosphorylation-dependent, direct interaction with both µ1 and µ2(17); µ2 is the medium subunit of AP-2, a clathrin adaptor that mediates endocytosis [reviewed in (19)]. We showed that basic regions 1 and 2 on µ1 were required for the interaction with the Vpu CD, indicating a remarkable convergence in how two viral proteins, Nef and Vpu, co-opt AP-1(17). Moreover, we showed that the analogous basic regions on µ2 were required for the interaction with Vpu(17).

These results raised the question of whether the CDs of Furin and other cellular proteins that contain PSACs also bind directly to µ1 and µ2, and if so, whether they utilize the same basic regions as Nef and Vpu. Recent data support this notion by showing that the Furin PSAC binds to basic region 2 of µ1(20). Somewhat paradoxically, a µ1 protein in which this basic region is mutated failed to affect the trafficking of Furin, a result attributed to the presence of additional, potentially redundant sorting signals in the CD of the protein(20). Whether the Furin CD bound µ2 was not tested.

During our study of cellular multipass transmembrane proteins in the Serinc family, some of which inhibit the infectivity of retroviruses including HIV-1 and are counteracted by Nef(21,22), we noticed a potential PSAC of sequence SGASDEED in a cytoplasmic loop of Serinc3. Thus, we herein used Serinc3, Furin, and Vpu as prototypic as well as novel PSAC-containing viral and cellular proteins to test the hypothesis that PSACs bind to µ subunits at specific basic regions. We found that the CD of Furin bound to µ1 and µ2 in a phosphoylation-dependent manner, as did the putative PSAC-containing cytoplasmic loop of Serinc3. Whereas the interaction of the Furin CD with µ2 strictly required none of the five individual basic regions tested, the interaction of the Serinc3 loop with µ2 required a basic region previously reported to bind the acidic plasma membrane lipid phosphatidylinositol-4,5-bisphosphate (PIP2) [reviewed in (23)].

These data suggest that while PSACs bind µ subunits, the specific PSACs within different viral and cellular proteins recognize different basic regions on the µ surface. The physiologic relevance of this specificity is not yet clear, but it might provide the opportunity to inhibit the interaction of viral proteins such as Nef and Vpu with the µ subunits without inhibiting the interaction of at least some cellular proteins such as Furin and Serinc3. Finally, we show that the PSAC in Serinc3 has varied considerably during the evolution of placental mammals, specifically at the sites of the key serines, which appear to have co-evolved in a manner consistent with their joint contribution to µ-binding. Since such positive selection is often indicative of a viral-host conflict (reviewed in(24)], we speculate that the µ-binding activity of Serinc3 might have varied over the course of evolution in response to the presence or absence of retroviruses that, like HIV-1, encode Serinc-antagonists.

## Results

### Phosphorylation-dependent binding of the cytoplasmic domains of Furin and Serinc3 to the µ subunits of AP-1 and AP-2; Serinc3 contains a phosphoserine acidic cluster (PSAC) sorting signal

Based on our previous studies of the HIV-1 proteins Nef and Vpu and how these proteins interact with the µ subunits of heterotetrameric clathrin adaptors, we tested the general hypothesis that PSAC sorting motifs bind directly to the basic regions of µ subunits. To do this, we examined the cellular protein Furin, a type I transmembrane protein whose cytoplasmic domain (CD) contains the canonical PSAC motif SDSDEEDE(4), and Serinc3, a multi-pass transmembrane protein whose predicted long cytoplasmic loop (herein designated loop 10) contains a similar sequence SGASDEED. HIV-1 Vpu is a type I transmembrane protein whose PSAC sequence is EDSGNESE(17). The predicted topologies of these three proteins and the sequences of their PSAC-containing CDs are shown in Figure 1. Potential additional AP-binding motifs are also present in Vpu and Furin(17,25-27); these include the Yxxϕ sequences that have the potential to bind µ subunits via a well-defined “two prong in socket” mechanism(5). Serines that are either reported or shown herein (Figure 1C) to be phosphorylated by casein kinase II (CK II) are indicated(28,29).

**Figure 1.**
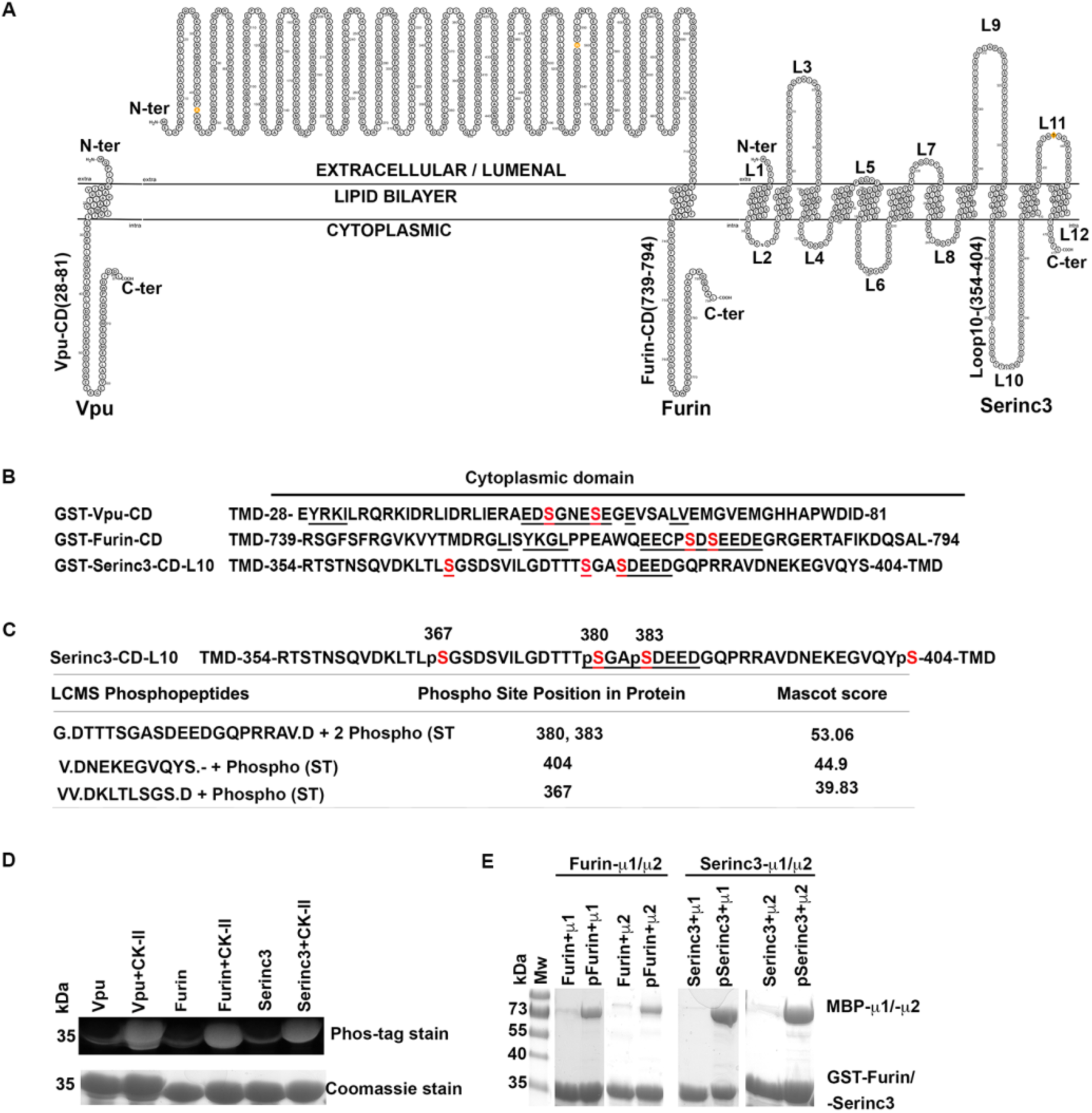
Cytoplasmic domains containing phosphoserine acidic clusters (PSACs) bind directly to the µ subunits of AP-1 and AP-2. **A.** Diagram of the transmembrane proteins HIV-1 Vpu, Furin, and Serinc3. Vpu and Furin are single-pass, type I transmembrane proteins. Serinc3 is a multi-pass transmembrane protein. Extracellular/luminal and cytoplasmic regions (loops) of Serinc3 are numbered N-terminal to C-terminal as L1-L12. L10 contains a potential PSAC. **B.** Cytoplasmic domains (CDs) of Vpu and Furin, and L10 of Serinc3. TMD; transmembrane domain. Potential clathrin adaptor protein (AP) complex binding motifs, including PSACs, are underlined. Serines in red are known or potential sites of phosphorylation by casein kinase II (CK-II). **C.** Sites of serine phosphorylation in L10 of Serinc3. GST-Serinc3-L10 was co-expressed with CK-II in *E.coli*, purified, and analyzed by liquid chromatography/mass spectrometry (LC/MS). Phospho-peptides and serine-threonine (ST) phosphorylation are indicated, as are the sites of phosphorylation. Mascot scores for the peptide matches are indicated. **D.** Phosphorylation of the CDs of Vpu and Furin, and L10 of Serinc3 detected by Phos-tag staining. GST-Vpu-CD, GST-Furin-CD, or GST-Serinc3-L10 was expressed in *E.coli* either with or without co-expression of CK-II. The proteins were purified by affinity chromatography using glutathione-sepharose, ion exchange and size-exclusion chromatography, and then stained either with Phos-tag stain or Coomassie blue. **E.** The CD of Furin and L10 of Serinc3 bind µ subunits in a CK-II-dependent manner. GST-Furin-CD or GST-Serinc3-L10 were expressed in *E.coli* either with (“p”) or without (no “p”) co-expression of CK-II. The purified proteins were then used to capture the C-terminal domains of either µ1 or µ2, each fused to maltose binding protein (MBP) as a solubility tag. SDS/PAGE gels were stained with Coomassie blue. Phospho-Furin CD and phospho-Serinc3-L10 each bound both µ1 and µ2.

We co-expressed each of these CDs as GST-fusion proteins in *E.coli* together with CKII. We analyzed phospho-GST-Serinc3 loop 10 by LC/MS and identified several peptides that together revealed CKII-mediated phosphorylation at positions 367, 380, 383, and 404 (Figure 1C); serines 380 and 383 are part of the putative PSAC. We analyzed all the GST-CD fusion proteins (GST-Serinc3 loop 10, GST-Vpu-CD, and GST-Furin-CD) by Phos-tag stain, and each was phosphorylated by CKII (Figure 1D).

To study the interactions of these phosphorylated CDs with the µ subunits, we used fusions of the C-terminal two-thirds of µ1 or µ2 to maltose binding protein (MBP; a solubility-enhancing tag) in pulldown assays in which the GST-CD fusion proteins were the bait (Figure 1E). Similar to our findings using the same assay to study the CD of Vpu, the CD of Furin and loop 10 of Serinc3 each bound to µ1 and µ2 in a CKII-dependent manner, which presumably reflected the requirement for serine- or threonine-phosphorylation.

### Each of the two phosphoserines of the Serinc3 PSAC contributes to binding, and the major contributor, phosphoserine 383, contributes partly via its negative charge

We next used the GST pulldown assay to identify the requirements within Serinc3 loop 10 for binding to the μ subunits (Figure 2). These requirements were similar for μ1 and μ2 (compare Figure 2A and 2B). Substitution of both S380 and S383 with asparagines reduced μ-binding to the residual levels of the non-phosphorylated protein. Serines 380 and 383 were each individually important, but S383 was the more critical residue for binding to µ2. Binding to µ2 was substantial when S383 was substituted with aspartate, but only when the protein was phosphorylated (Figure 2B); these observations suggested that while negative charge is a key attribute required for binding, residues in addition to S383, such as S380, likely require phosphorylation for the loop to bind µ2. Notably, the S383D mutant bound µ1 relatively poorly, even when the protein was phosphorylated (Figure 2A). Figure 2C shows a reversed pulldown design in which the C-terminal two-thirds of µ2 are used as the bait; the data confirm that S380 and S383 in loop 10 are required for the interaction and further show that serines 356, 359, 367, and 371 are dispensable.

**Figure 2.**
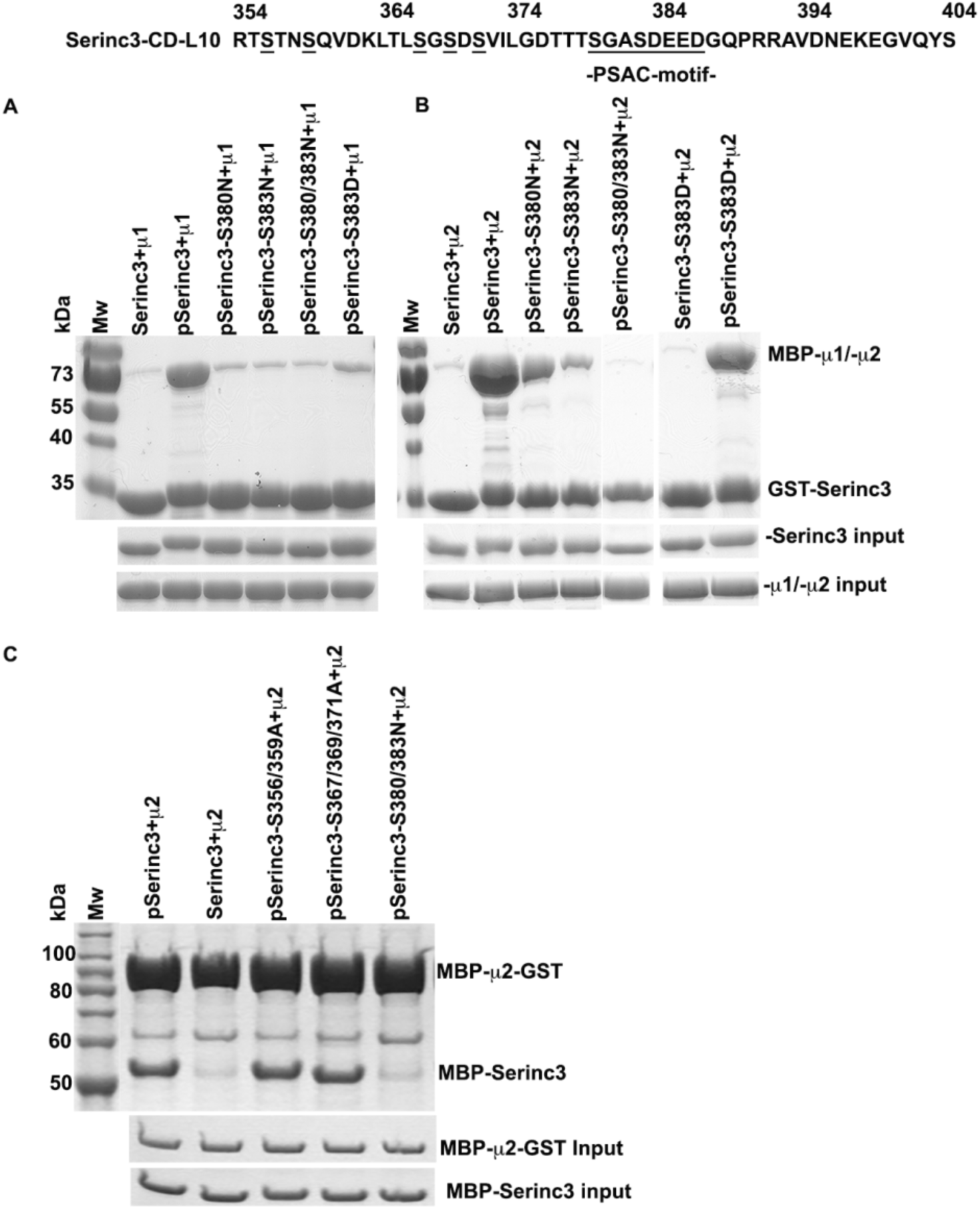
Mutational analysis of the PSAC of serinc3-L10 with respect to µ-binding. **A.** GST-Serinc3-L10 or the indicated S/N or S/D substitution mutants were expressed in *E.coli* either with (“p”) or without (no “p”) co-expression of CK-II; the purified proteins were used to pulldown MBP-µ1. B. Same experiment as in panel A except that the indicated proteins were used to pulldown MBP-µ2. The maximal interaction between Serinc3-L10 with the µ subunits required co-expression of CK-II and both serines S380 and S383. C. MBP-µ2-GST was used to pulldown phospho-MBP-Serinc3-L10. This “reverse” pulldown relative to panels A and B confirmed the key roles of S380 and S383 in the interaction, and it indicated that S356, S359, S367,369 and S371 are dispensable.

### The cytoplasmic domains of Vpu, Furin, or Serinc3 cytoplasmic loop 10 form stable complexes with µ2 at low or sub-micromolar affinity

We next sought to confirm that the interactions between the CDs of Vpu, Furin, and Serinc 3 loop 10 with μ2 were stable and to define their kinetics and affinities. We used size exclusion chromatography (SEC) to analyze equimolar mixtures of the GST-fusion proteins containing the CDs of Vpu or Furin or Serinc3 loop 10 and MBP-µ2 (Figure 3A). The data indicate that each of these CDs forms stable complexes with µ2, indicated in each case by the peak in the elution profile designated P1.

**Figure 3.**
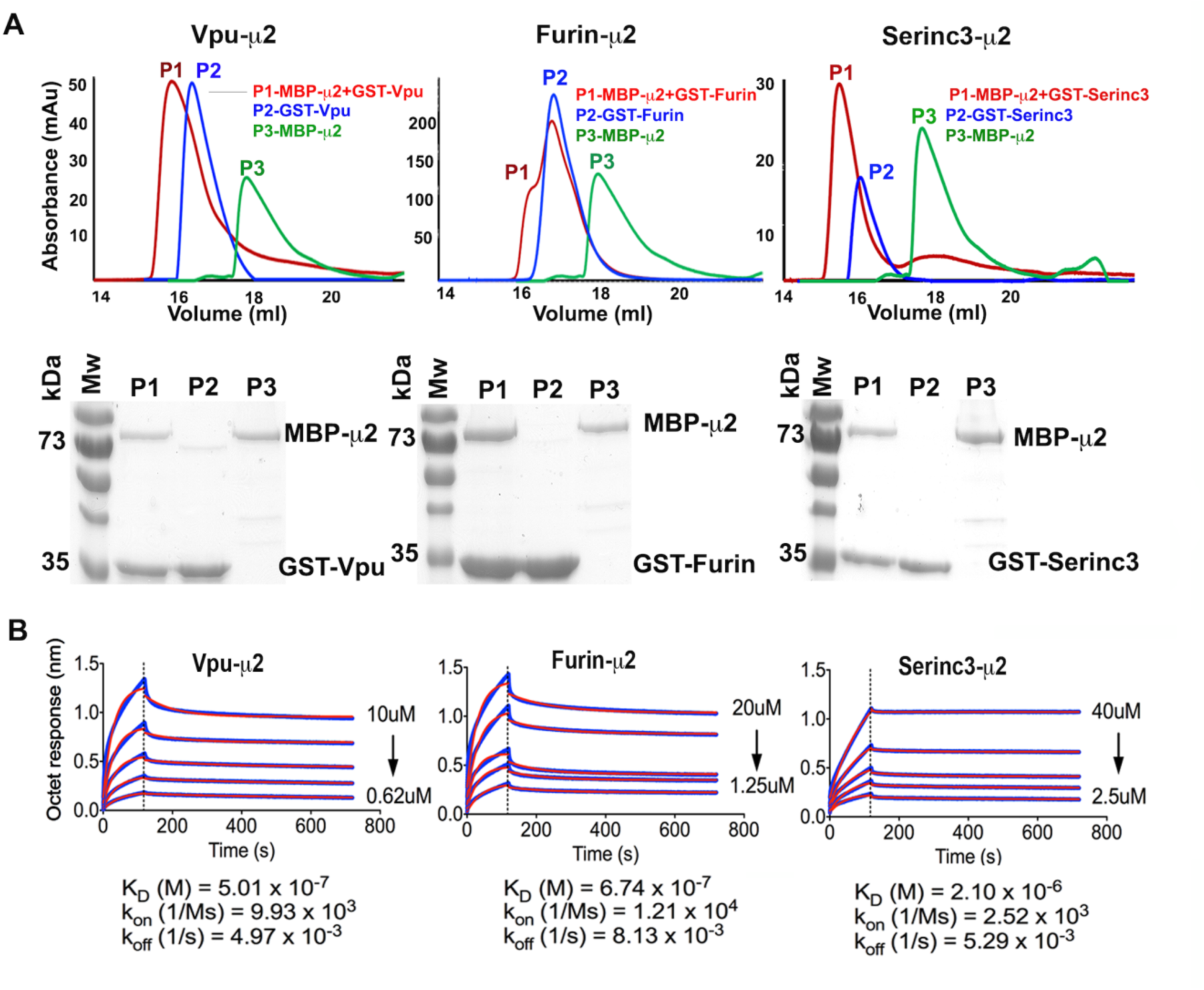
Binding between the Vpu-CD, Furin-CD or loop 10 of Serinc3 and µ2 measured by size exclusion chromatography (SEC) and biolayer interferometry. **A.** GST-Vpu-CD, GST-Furin-CD and GST-Serinc3L10 were fractionated by SEC either alone (P2, blue) or after mixing in equimolar amounts with MBP-µ2 (P1; red). MBP-µ2 was also fractionated alone (P3; green). Top: protein absorbance values vs. elution volume for the three independent SEC experiments. Bottom: Coomassie blue-stained SDS/PAGE gels of the indicated peaks (P1, P2, P3); P1 contains the protein-complexes. **B.** Biolayer interferometry. His-tagged MBP-µ2 was immobilized on a Ni-NTA chip surface and GST-Vpu-CD, GST-Furin-CD, and GST-Serinc3-L10 were used as the analytes at the indicated concentrations. The vertical dotted lines separate the association (left) from the dissociation (right) steps of the experiments. The dissociation constants (K_D_) between µ2 and different PSAC-containing CDs are shown along with the on- and off-rates.

To define the kinetics and affinities of these interactions, we used biolayer interferometry. We immobilized poly-histidine-tagged MBP-µ2 on a Ni-NTA chip surface and used the GST-CD fusion proteins at different concentrations as the analytes. The association and dissociation steps of the experiments are shown in Figure 3B, along with the derived on/off-rates and the K_D_ value for each interaction. These data indicated that on-rates of the Vpu-µ2 and Furin-µ2 interactions are similar to each other and faster than that of Serinc3 loop 10-µ2, whereas the off-rates of all three interactions were similar. The K_D_ values of the Vpu-µ2 and Furin-µ2 interactions are sub-micromolar, whereas that of the Serinc3 loop 10-µ2 interaction is low micromolar. The K_D_ of the Furin CD-µ2 interaction, determined here by biolayer interferometry, was 0.67 µM, substantially less than the published K_D_ of the interaction of a phosphorylated Furin peptide with µ1 determined by isothermal calorimetry (22 µM) or surface plasmon resonance (35 µM)(20). This could reflect differences between µ1 and µ2; the use of a relatively short peptide in the studies of µ1 rather than the entire CD as used here; or the presence of GST-tags in our analyte proteins, which could stimulate dimerization and consequently increase the apparent affinity of the CDs for µ2 in the above experiments.

### Basic domains on the µ subunits: Serinc3 loop 10 requires basic region 4 for binding, a region previously reported to bind PIP2

We next sought to map the basic regions on the surface of the µ2 subunit that contribute to the binding of these PSAC-containing CDs. Figure 4A shows a surface representation of µ1(basic regions in blue) complexed to a fusion protein consisting of the CD of the MHC-I α-chain fused to HIV-1 Nef (red ribbon and red space-fill). This previously published crystal structure led to our identification of two basic regions on µ1, herein designated 1 and 2, that are required for the interaction with MHC-I-Nef(12). Some but not all of the charged lysine and arginine residues in these regions are conserved (see Figure 4D, which shows a structure-based alignment of µ1 and µ2). The analogous basic regions (1 and 2) are indicated on a surface representation of µ2 (Figure 4B); note that basic region 1 of µ2 contains a lysine analogous to arginine 225 of µ1, but it is an unstructured region not shown in Figure 4B. Also note that basic region 2 of µ2 consists of only two lysine residues (K308 and K312); residues analogous to K274 and R303 of µ1 are not present. We recently showed that basic regions 1 and 2 are each required for the binding of the Vpu CD to either µ1 or µ2(17). We extended this analysis to the interactions of µ2 with the Furin CD and loop 10 of Serinc3 (Figure 4E): the pulldown data indicate that neither basic region 1 nor 2 of µ2 is required for the interactions, although basic region 2 might be contributing given the slightly reduced signal.

**Figure 4.**
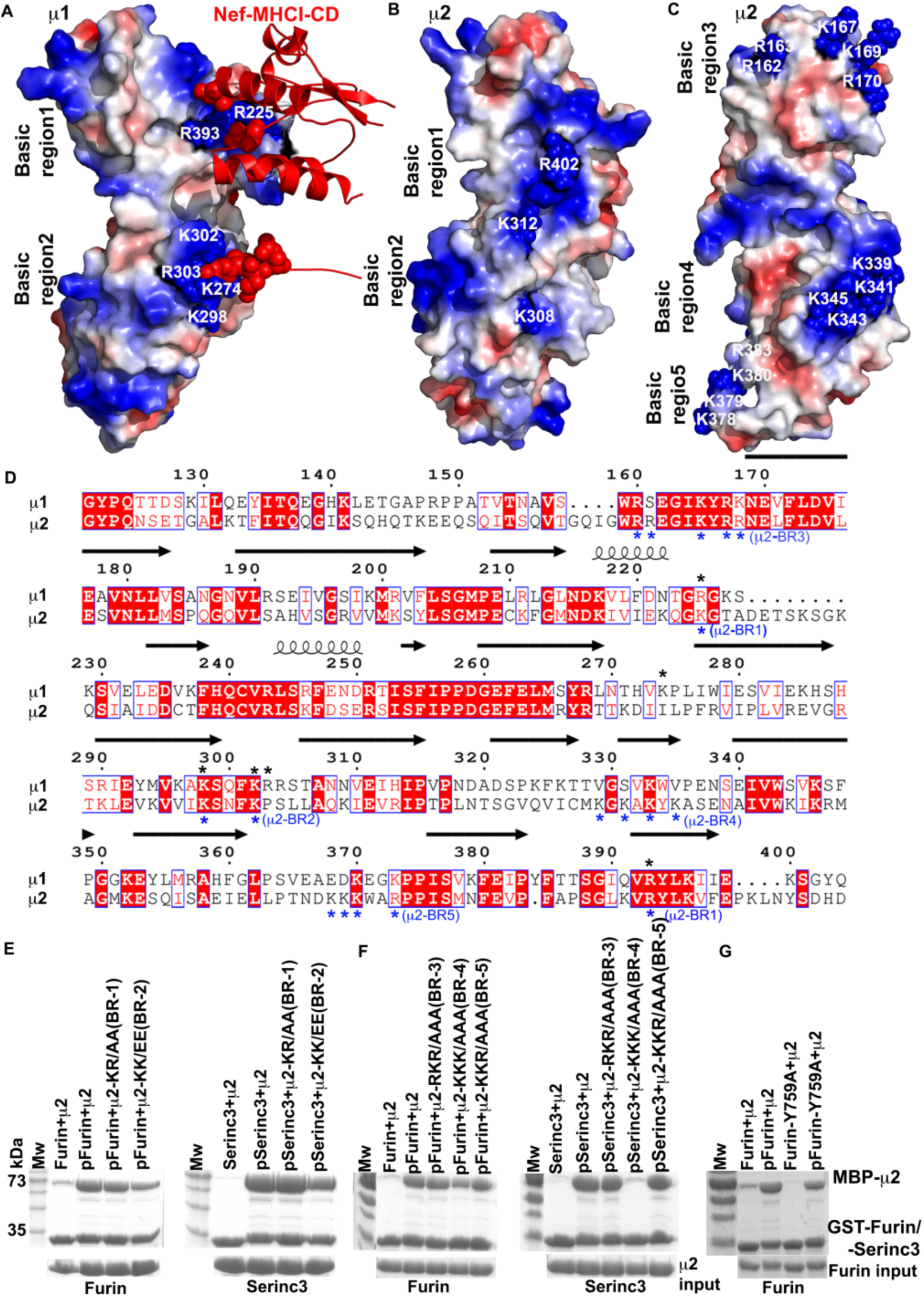
Identification of basic patches on µ2 and their roles in binding the CD of Furin and loop 10 of Serinc3. **A.** Surface representation of the C-terminal two-thirds of µ1 complexed with HIV-1 Nef fused to the CD of the class I MHC α chain (red ribbon; red space-fill for the Nef acidic cluster) (PDB code:4EN2). Basic regions (blue) 1 and 2 (numbered residues) were shown previously to be essential for the interaction. **B**. Surface representation of µ2 (PDB code:1BW8) with basic regions analogous to regions 1 and 2 of µ1. **C.** Surface representation of µ2 rotated relative to panel B and indicating additional basic regions 3, 4, and 5. Region 4 is a binding site for the acidic membrane lipid PIP2. **D.** Structure-based sequence alignment of µ1 and µ2 was done using ESPript3(49). Black asterisks indicate the positions of residues constituting basic regions 1 and 2 of µ1. Blue asterisks indicate basic regions 1-5 of µ2. Secondary structures are indicated by arrows (β-stands) and coils (α-helices) above the sequence. Red boxes indicate strict identity. **E.** Pulldown of MBP-µ2 or the indicated basic region (“BR”) mutants by GST-Furin-CD or GST-Serinc3-L10. “p” indicates co-expression of the GST-fusion proteins with CK-II. Neither basic region 1 nor 2 of µ2 is required for binding. **F.** Pulldown of MBP-µ2 or the indicated basic region mutants by GST-Furin-CD or GST-Serinc3-L10. Basic region 4 (BR-4) is required for binding serinc3-L10. **G.** Pulldown of MBP-µ2 by GST-Furin-CD or a YxxΦ-mutant of the Furin CD (Y759A). Y759 is required for the phosphorylation-independent binding of the Furin-CD to µ2 but contributes minimally to the binding of the phosphorylated Furin-CD.

Given these results, we defined and mutated three additional basic regions on the surface of µ2, designated 3, 4, and 5 and shown in Figure 4C in a surface representation of µ2 that is rotated relative to the view shown in Figure 4B. The roles of these basic regions in the interactions of µ2 with the Furin CD and loop 10 of Serinc3 were tested in pulldown assays (Figure 4F): the data indicated that basic regions 3 and 5 are dispensable for these interactions, whereas basic region 4 is required for the binding of Serinc3 loop 10 and might contribute slightly to the binding of the Furin CD. Overall, these data indicate that the Furin CD and loop 10 of Serinc3 bind µ2 with requirements for basic regions on the µ-surface different from those we previously defined for Vpu. While the binding of Furin persisted despite mutation of any of five different basic regions, binding of loop 10 of Serinc3 required basic region 4, which has been reported previously as a binding site for the plasma membrane specific phospholipid PIP2(30,31).

Since none of the five basic regions on µ2 were critical for interacting with the Furin, we tested the possibility that residual binding was due to the Yxxφ sequence (Y_759_KGL) upstream of the Furin PSAC (see Figure 1B)(8). Figure 4G shows that while the residual binding of the Furin CD to µ2 in the absence of phosphorylation was dependent on Y759, the Y759A substitution only minimally decreased binding to µ2 when the Furin CD was phosphorylated. These data indicate that the persistent binding of the wild type phosphorylated Furin CD to the single-basic-region µ-mutants shown in Figure 4E and 4F is not attributable to the activity of the Yxxϕ sequence.

### Genetic variation in the Serinc3 PSAC suggests the possibility of host-pathogen conflict

We previously noted that the sites of S380 and S383 in Serinc3 are under positive (diversifying) selection among primates(32). In some cases, the 383 position is substituted with asparagine, which as shown in Figure 2 is highly detrimental to µ-binding. Here, we examined more completely how this position has varied among placental mammals. A phylogeny of eutheria based on *serinc3* mRNA sequence is shown in Figure 5; the branches with amino acid substitutions at position 380 and 383 are color-coded (blue: 380; red: 383; purple: 380 and 383). Position 383 of Serinc3 has alternated between serine, asparagine, and glycine during eutherian evolution. Of the 36 non-synonymous substitutions we detected at these sites across the phylogeny, 32 involved substitutions to or from a serine (Figure 5). Positions 380 and 383 tend to co-evolve (probability = 0.999 in the Bayesian graphical model), favoring a model in which the amino acid at site 383 is conditionally dependent on the amino acid at site 380. In general, position 380 has evolved towards serine, whereas position 383 has repeatedly evolved away from serine. Nonetheless, the phylogeny indicates four acquisitions of serine at position 383. These results are consistent with the joint contribution of the two serines to µ-binding and indicate that the ability of loop 10 of Serinc3 to bind µ subunits has waxed and waned among placental mammals. This variation could relate to some form of selective pressure, conceivably that of retroviral proteins that counteract the ability of Serinc3 to inhibit infectivity.

**Figure 5.**
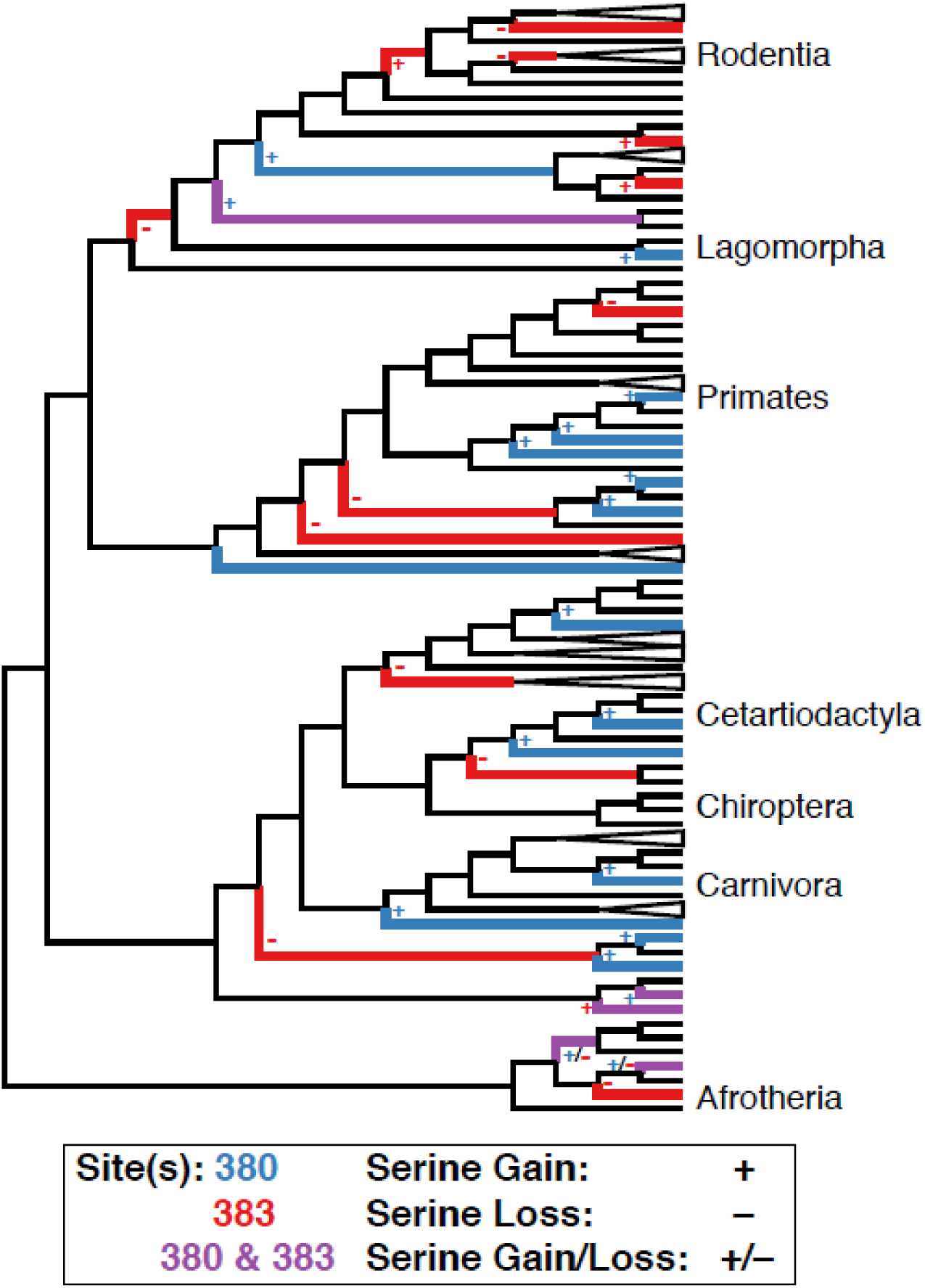
Cladogram of placental mammals depicting the coordinated evolution at codon position 380 and 383 of the Serinc3 loop 10. Blue branches indicated amino acid substitutions at site 380, red branches indicate amino acid substitutions at 383, and purple branches indicate amino acid substitutions at both sites. Pluses indicate a substitution toward serine; minuses indicate a substitution away from serine. The cladogram is based on a maximum likelihood tree of 133 eutherian *serinc3* mRNA sequences. Major mammalian clades are identified.

### The Serinc3 PSAC appears dispensable for the inhibition of HIV-1 infectivity and for counteraction by Nef

Given the forgoing observations, we tested whether the PSAC of Serinc3 affects either its activity in inhibiting viral infectivity or its susceptibility to counteraction by the HIV-1 Nef protein. Insofar as exemplified by Serinc5, Nef reduces the amount of Serinc proteins in virions in a clathrin- and AP-2-dependent manner, thereby counteracting the negative-effect of the Serincs on infectivity(21,22). We measured the infectivity of HIV-1 virions, either genetically *nef*-positive (wild type) or -negative (ΔNef), produced either in the presence or absence of wild type Serinc3 or three different PSAC-mutants: a deletion of the sequence SDEED, an alanine substitution mutant of S383, or an alanine substitution mutant of the DEED acidic sequence. Each of the Serinc PSAC mutants inhibited the infectivity of virions produced in the absence of Nef, and each was similarly susceptible to antagonism by Nef (Figure 6A). Western blot indicated that all the Serinc3 proteins were detected in partially purified preparations of virions, the apparent site of action of the Serincs(21). When compared to the wild type protein, the Serinc3 mutant in which the SDEED sequence was deleted and the mutant in which the DEED sequence was replaced with alanines, but not the S383A mutant, were relatively better expressed at steady-state in cells and accumulated more efficiently in virions. This observation leads to the hypothesis that the acidic residues lead to endo-lysosomal degradation. Nonetheless, each of the Serinc3 proteins was excluded from virions by Nef to similar extents. Overall, these data indicated that the serinc3 PSAC is not a substantial determinant of antiretroviral activity or of the susceptibility of the protein to Nef-mediated antagonism.

**Figure 6.**
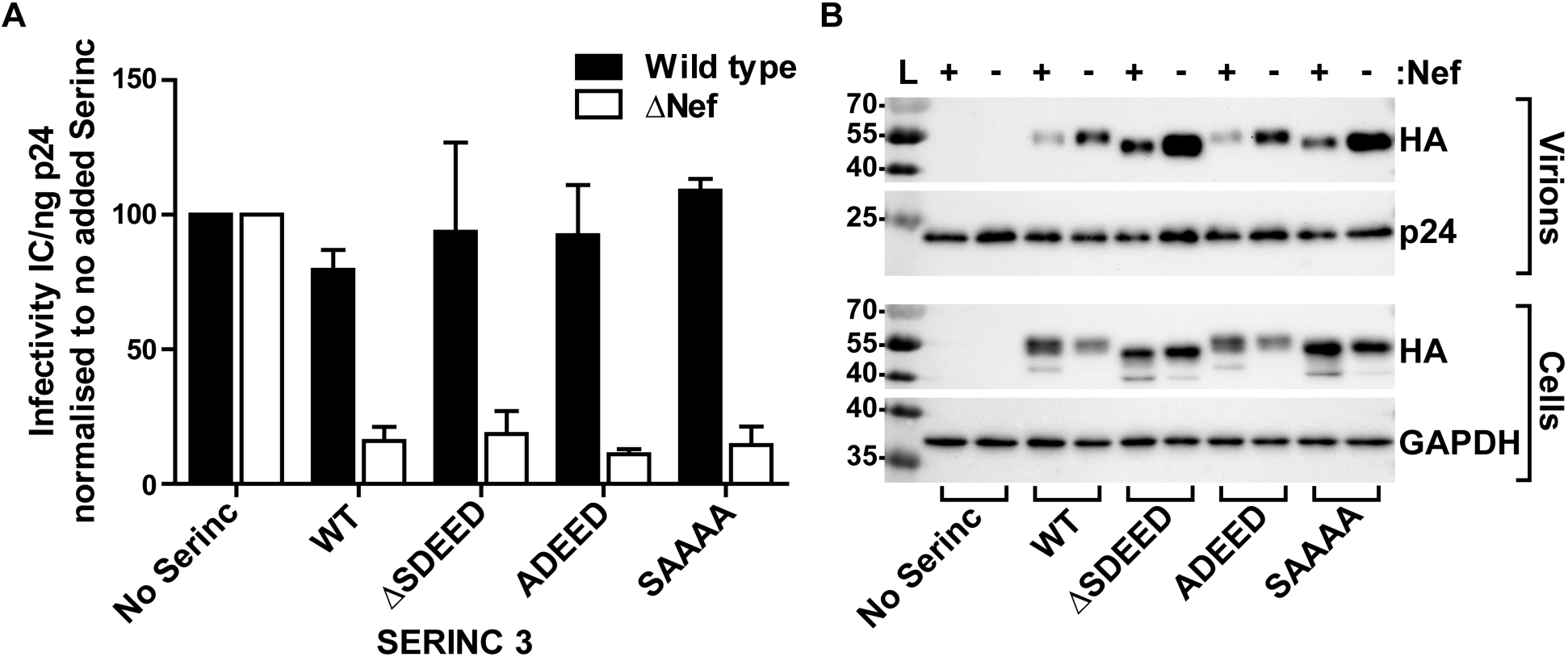
The serinc3 PSAC is dispensable for antiviral activity and for counteraction by HIV-1 Nef. **A.** Relative infectivity of HIV-1 virions, either wild type of *nef*-negative (ΔNef) when produced from Jurkat T cells whose ORFs for *serinc3* and *serinc5* were disrupted by CRISPR-editing. The indicated Serinc3 proteins were expressed by transfection of the cells with plasmids. The SDEED sequence of the PSAC was either deleted or partially substituted with alanines as indicated. Viral infectivity was measured on HeLa-CD4 indicator cells; the infectivity (initially calculated as infectious centers (IC) per ng of p24 capsid antigen) of each virus (wild type or *nef*-negative) in the presence of Serinc3 is expressed relative to its no-Serinc control (set at 100). Error bars are plus/minus the standard deviation of triplicate, independent experiments. **B.** Western blots of a representative experiment showing Serinc3 (HA) in virions (top) and virion-producer cells (bottom). p24 is the virion-capsid antigen; GAPDH is a cellular protein used as a loading control.

**Figure 7.**
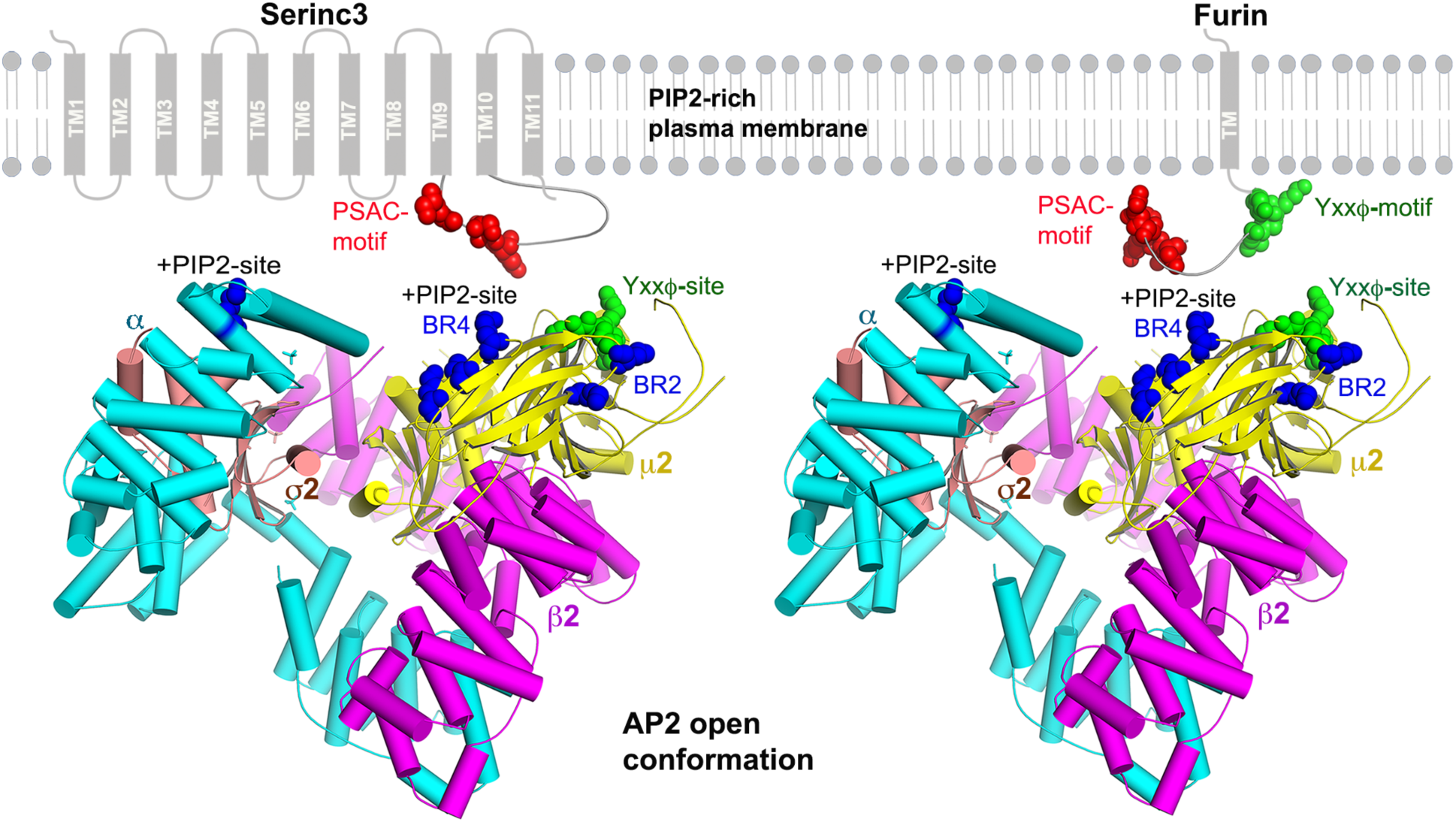
Possible mode of interaction of AP-2 with the PSAC-containing CDs of Serinc3 and Furin. The AP2-core is shown in an “open” conformation (PDB code:2XA7)(23). A peptide derived from TGN38 and bound to the Yxxϕ-binding pocket of µ2 is shown by green spheres. Basic regions −2 and −4 are shown by blue spheres. Two PIP2-binding sites on AP-2 are indicated: one on the α subunit and the other on µ2, which is the same as basic region 4. The PIP2 sites and cargo-binding sites are co-planar and face the membrane. Serinc3 and Furin are shown schematically, with the TMs of Serinc3 numbered; loop 10, which contains the PSAC, is between TM9 and TM10. PSACs are red; Yxxϕ motifs are green. The distances from residue D176 of the Yxxϕ-binding site to the residues of basic region 4 are 26.A (to K341), 27.3A (to K343), and 30.9A (to K345); the distances from D176 to the residues of basic region 2 are 31.8A (to K308) and 20.3A (to K312). These distances are consistent with simultaneous binding of the Furin CD to the Yxxϕ-binding site via its Yxxϕ-motif and to basic region 4 or 2 via its PSAC.

## Discussion

We tested the hypothesis that the phosphoserine acidic cluster (PSAC) sorting signals found in the cytoplasmic domains (CDs) of viral and cellular proteins, here exemplified by HIV-1 Vpu, Furin, and Serinc3, bind to specific basic regions on the µ subunits of the clathrin adaptor protein complexes. This hypothesis was initially derived from our identification of two basic regions on µ1 (the µ subunit of AP-1) that interact with acidic residues within the HIV-1 Nef and MHC-I α chain cytoplasmic domain complex(12). It was reinforced by our more recent observation that these basic regions on µ1, and the analogous basic regions on µ2 (the µ subunit of AP-2), are required for interaction with the cytoplasmic domain of the HIV-1 immuno-modulatory protein Vpu(17). Thus, we proposed that Vpu contains a PSAC similar to that of the prototypical sequence found in the cellular protein Furin(4), and that the cytoplasmic domain of Furin might similarly bind the µ subunits via its PSAC.

Here, we found that the CD of Furin binds µ1 and µ2 in a phosphorylation-dependent manner. Moreover, we identified a novel PSAC in a cytoplasmic loop of the cellular protein Serinc3, which restricts retroviral infectivity(22). The loop of Serinc3 binds to both µ1 and µ2. We documented that each of the serines in the PSAC of Serinc3 are phosphorylated by casein kinase II in vitro, and each contributes to binding to both µ1 and µ2. While the CDs of Vpu, Furin, and Serinc3 each bind to µ2 with low or sub-micromolar affinity, they appear to interact with different basic regions on the surface of µ2. Remarkably, neither Furin nor Serinc3 required either of the basic regions that are utilized by Vpu(17). Serinc3 specifically required a basic region on µ2 that has previously been implicated as a binding site for the acidic phospholipid PIP2(23,30,31). Moreover, of five basic regions tested on µ2, none were individually required for the interaction with the phosphorylated cytoplasmic domain of Furin. These data suggest the possibility of redundancy in the use of specific basic regions by Furin, and non-overlapping specificity in the sites recognized by other PSAC-containing CDs such as those of Vpu and Serinc3.

What could be the biological relevance of multiple binding sites for PSACs on the surface of µ? These multiple sites might provide flexibility for how the PSAC motifs in various CDs can avail themselves of high-affinity binding. For example, this flexibility might allow the PSACs in different proteins to function while at different distances from the membrane, or it might allow the simultaneous use of different types of AP-binding signals within a given protein’s CD. Both Furin and Vpu, for example, encode potential Yxxϕ and ExxxLϕ signals at various distances from their PSACs(8,17,26,27). The binding site for Yxxϕ signals is, like those for the PSAC signals, on the µ subunits(5). The concomitant recognition of Yxxϕ and PSAC signals in the same CD by the µ subunit is plausibly exemplified by the HIV-1 Nef/MHC-I α chain CD/µ1 complex, in which a tyrosine in the MHC-I CD acts as an partial Yxxϕ signal and occupies the canonical binding pocket on µ1, while acidic residues in Nef and the MHC-I CD (although not PSACs per se) simultaneously bind basic regions on the surface of µ1(12). The tyrosine of the MHC-I CD (Y320) is only seven residues upstream of the aspartate (D327) that binds basic region 1. By analogy, the CD of Furin contains a Yxxϕ signal whose tyrosine is 10 residues upstream of the closest glutamic acid residue that could contribute to the PSAC; nonetheless, the Yxxϕ motif in the Furin CD contributed relatively little to the interaction under the conditions tested here, and basic region 1 of μ2 was dispensable. Instead, basic region 2 appeared to contribute modestly to the binding of the Furin CD to µ2, a result potentially consistent with the more substantial role reported for this region in binding to µ1 as discussed below(20). Basic region 4 also appeared to contribute to the binding of the Furin CD to µ2, but its role in binding loop 10 of Serinc3 was much more striking.

How can the Serinc3 PSAC (and perhaps the PSAC in the Furin CD) use the basic region on µ2 that has been associated previously with the binding of PIP2(30,31)? Notably, the interaction of AP-2 with PIP2 is an apparent mechanism by which AP-2 is specifically recruited to the plasma membrane rather than to internal endosomal membranes. At least two binding sites for PIP2 on AP-2 have been described: one on the µ subunit (herein basic region 4) and another on the larger α subunit(23,30,31). These sites have been proposed to act sequentially, with the site on the α subunit mediating an initial low-affinity interaction that is subsequently stabilized by a conformational change in the complex to an “open” form (partly due to phosphorylation of µ2 itself)(33) that exposes the C-terminal two-thirds of µ2 and allows it to bind Yxxϕ signals in transmembrane protein cargo and another molecule of PIP2. Conceivably, the Serinc3 PSAC might partly substitute for PIP2 in this scenario, stabilizing the interaction between AP-2, the lipid bilayer, and the transmembrane protein cargo, here Serinc3 itself. Figure7 depicts a schematic representation of how the PSACs in Serinc3 and Furin could interact with AP-2, which is shown in an open conformation with a TGN38-derived peptide bound to µ2 at the Yxxϕ-binding site (23)

We previously reported that the sites in *serinc3* at which the human gene encodes serines of the PSAC in loop 10 (positions 380 and 383) are under positive (or diversifying) selection when non-human primate nucleotide sequences are compared(32). This genetic phenomenon is often the signature of a host-pathogen conflict, in which the host antiviral gene is subject to selection pressure to escape the ability of viral proteins to antagonize it(24). Thus, we remain intrigued that the ability of Serinc3 to bind µ2 via the PSAC in loop 10 appears to wax and wane over the phylogeny of placental mammals. While the Nef protein of HIV-1 antagonizes the inhibitory effects of Serinc family proteins on viral infectivity(21,22), this function is provided by the transmembrane protein glycoGag in murine leukemia virus (an oncogenic retrovirus) and by the S2 protein in equine infectious anemia virus (a lentivirus of horses)(34,35). In neither the mouse nor the horse is the PSAC signal intact: in the mouse the serines are replaced by asparagines and in the horse by glycines. Since these viral antagonists of the Serinc proteins presumably function by affecting membrane trafficking, whether the presence or absence of an intact PSAC motif in Serinc proteins affects their activity remains an interesting and open question. So far, we have shown only that the presence or absence of an intact PSAC has no influence on the activity of a single HIV-1 Nef protein as an antagonist of Serinc3, even though this cytoplasmic loop has recently been shown to be a determinant of the susceptibility of Serincs to Nef proteins(36).

Recently, a haploid cell genetic screen using a gene-disrupting retrovirus and a fusion protein of CD8 and the CD of Furin (mutated to lack its Yxxϕ signal) as a flow cytometric indicator revealed that µ1 is a target of the PSAC in Furin(20), a conclusion fully consistent with the data herein. These investigators also reported that basic region 2 of µ1, the site that interacts with the acidic cluster of Nef, is required for the interaction of the Furin CD with µ1 in vitro; this result is potentially but not necessarily at odds with the data herein, in which the analogous region of µ2 appears dispensable-although modestly contributory - for binding to the Furin CD. Nonetheless, the authors found that while a µ1 mutant in which basic region 2 was disrupted did not fully support the modulation of class I MHC by Nef, it paradoxically fully supported the trafficking of Furin; that is, basic region 2 of µ1 was dispensable for Furin-trafficking(20). Conceivably, the presence of additional AP binding signals in the CD of Furin could resolve this paradox, leading to the authors’ notion that redundancies in the sorting signals within the CDs of cellular proteins might render them less susceptible to disruption than the single binding sites in viral proteins(20). Another possibility, supported by the data herein, is that the interaction of the Furin CD with μ2 is a key determinant of Furin-trafficking.

Along a distinct line of reasoning, our data suggest that multiple basic regions on the µ subunits can be utilized differentially by the PSACs of different viral and cellular proteins. This scenario offers the potential to inhibit the interaction of viral proteins such as Nef and Vpu with specific basic regions on the µ subunits (namely, regions 1 and 2, herein), while leaving the trafficking of at least some cellular proteins such as Furin and Serinc3 relatively unaffected. Exactly how that might be accomplished awaits a high-resolution structural definition of these interactions.

## Methods and Materials

### Plasmids and human cell lines

The HeLa-derived reporter cell line P4.R5 (37) was obtained through the NIH AIDS Reagent Program, Division of AIDS, NIAID, NIH, from Nathaniel Landau. The cells were maintained in Dulbecco’s modified Eagle medium (DMEM) supplemented with 10 % fetal bovine serum (FBS), penicillin/streptomycin, and 1 µg/ml puromycin. The double-knockout Jurkat TAg T lymphoid cell line in which the *SERINC3* and *SERINC5* ORFs are disrupted was a kind gift from Heinrich Gottlinger (22) and was maintained in RPMI medium supplemented with 10% FBS and penicillin/streptomycin.

The proviral plasmids pNL4-3 and pNL4-3ΔNef have been described previously(38-40). The expression plasmid encoding a C-terminally HA epitope-tagged SERINC3 in the modified mammalian expression vector pBJ6 was a kind gift from Massimo Pizzato (21). The SERINC3-HA coding sequence was transferred to a pBJ5 expression vector by restriction digestion and ligation using NotI and EcoRI sites. Expression plasmids encoding pBJ5-SERINC3 ΔSDEED, ADEED, and SAAAA were generated by site-directed mutagenesis using a QuikChange II site-directed mutagenesis kit (Agilent Technologies); primers were synthesized by Integrated DNA Technologies.

### Cloning, expression, and purification of recombinant proteins

The HIV1-Vpu, Furin and Serinc3 constructs were designed based on their known or predicted cytoplasmic domains. The CDs of Vpu (28-81) and Furin (739-794) were cloned into the pGEX4T1 vector, which added a Glutathione-S-transferase (GST)-tag at the N-terminus of the protein. We used computational predictions to identify the topology of Serinc3 and its putative extracellular and intracellular loops. Loop10 of Serinc3 (residues 354-404) is predicted to be cytoplasmic, and by sequence inspection it contains a potential PSAC motif: 383-SDEED-387, which represents a consensus Casein Kinase-II (CK-II) site. We cloned this loop into the pGEX4T1 vector. For the AP-1 and AP-2 medium subunits (µ1 and µ2), we used previously created(12) truncated versions of µ1(158-423) and µ2(159-435) in the pMAT9s vector, in which the μ sequences are fused to that of maltose binding protein (MBP). For reverse pulldown studies, we created MBP-µ2(159-435)-GST and MBP-Serinc3(355-405) constructs. The GST-tag was added to the C-terminus of a previously created MBP-μ2CTD construct with a tobacco etch virus (TEV) protease cut-site between μ2CTD and GST in the pMAT9s vector. MBP-SERINC3 (355-405) constructs were prepared similarly with a SARS M^pro^-(41,42)cleaveable 6xHis tag at the C-terminus in the pMAT9s vector; these constructs were cloned using ligation-independent Gibson assembly.

The GST-tagged Vpu, Furin, Serinc3-Loop10 and MBP tagged Serinc3-Loop10 proteins were expressed in BL21(DE3) competent cells (New England Biolabs). Protein expression was induced with 0.1 mM isopropyl β-d-thiogalactopyranoside (IPTG) at OD600 of 0.6∼0.8 at 16°C overnight. To make phosphorylated proteins, the GST-fusions were co-expressed with the α and β subunits of Casein Kinase II (CK-II) cloned into the pCDFDuet vector. MBP-µ1, MBP-µ2 and MBP-µ2-GST proteins were co-expressed with the pGro7 chaperone vector(43,44) and additionally induced with 1.5 g/L media L-(+)-arabinose at OD600 ∼ 0.2. Cell pellets were resuspended in appropriate binding buffers and lysed either by French press homogenizer or by micofluidization. Lysates were clarified by centrifugation at 14,000 rpm. GST-Vpu, GST-Furin and GST-Serinc3 were purified by GST-affinity, HiPrep Q anion exchange, and S200pg chromatography. MBP-µ1 and MBP-µ2 proteins were purified by His-select Ni-affinity gel, HiPrep S cation exchange, and S200pg chromatography. MBP-μ2CTD-GST was purified using Ni-NTA agarose, followed by a GST affinity column and gel filtration chromatography on a Superdex 200 prep-grade column. MBP-SERINC3 lysate was supplemented with 10 mM NaF and purified by Ni-NTA, HiTrap Q anion exchange chromatography, and S200pg chromatography.

All mutations in Furin, Serinc3 and µ2 were generated using the Quick-change mutagenesis kit (Agilent Technologies, La Jolla, CA); mutant proteins were expressed and purified as described above.

### Analysis of phosphorylation by LC/MS

GST-Serinc3 protein was co-expressed with CK-II in BL21 (DE3) cells, and purified samples were analyzed by LC/MS as follows:

In solution protein digestion: samples (95 µg in 5.0µl Tris buffer including 2mM DTT) were denatured by the addition of 5.0 µl 8M urea, 0.4M ammonium bicarbonate. The proteins were reduced by the addition of 1.0µl 45mM dithiothreitol (Pierce Thermo Scientific) and incubation at 37°C for 30 minutes, and then alkylated with the addition of 1.25µl 100mM iodoacetamide (Sigma-Aldrich) with incubation in the dark at room temperature for 30 minutes. The urea concentration was adjusted to 2M by the addition of 7.75µl of water. Samples were then enzymatically digested with 2.0µg of trypsin (Promega Seq. Grade Mod. Trypsin) and incubation at 37°C for 16 hours. Samples were desalted using C18 MacroSpin columns (The Nest Group) following the manufacturer’s directions with peptides eluted with 0.1% TFA, 80% acetonitrile. Eluted samples were dried and dissolved in MS loading buffer (2% acetonitrile, 0.2% trifluoroacetic acid). A Nanodrop (Thermo Scientific Nanodrop 2000 UV-Vis Spectrophotometer) was used to measure the protein concentrations (A260/A280). An aliquot of each sample was then diluted with MS loading buffer to 0.02µg/µl, with 0.1µg (5 µl) injected for LC-MS/MS analysis.

LC-MS/MS: LC-MS/MS analysis was performed on a Thermo Scientific Q Exactive Plus equipped with a Waters nanoAcquity UPLC system utilizing a binary solvent system (Buffer A: 100% water, 0.1% formic acid; Buffer B: 100% acetonitrile, 0.1% formic acid). Trapping was performed at 5µl/min, 97% Buffer A for 3 min using a Waters Symmetry® C18 180µm x 20mm trap column. Peptides were separated using an ACQUITY UPLC PST (BEH) C18 nanoACQUITY Column 1.7 µm, 75 µm x 250 mm (37°C) and eluted at 300 nl/min with the following gradient: 3% buffer B at initial conditions; 5% B at 1 minute; 35% B at 50 minutes; 50% B at 60 minutes; 90% B at 65 minutes; 90% B at 70 min; return to initial conditions at 71 minutes. MS was acquired in profile mode over the 300-1,700 m/z range using 1 microscan, 70,000 resolution, AGC target of 3E6, and a maximum injection time of 45 ms. Data dependent MS/MS were acquired in centroid mode on the top 20 precursors per MS scan using 1 microscan, 17,500 resolution, AGC target of 1E5, maximum injection time of 100 ms, and an isolation window of 1.7 m/z. Precursors were fragmented by HCD activation with a collision energy of 28%. MS/MS were collected on species with an intensity threshold of 2E4, charge states 2-6, and peptide match preferred. Dynamic exclusion was set to 20 seconds.

Peptide identification: data was analyzed using Proteome Discoverer software (version 1.3, Thermo Scientific) and searched in-house using the Mascot algorithm (version 2.6.0) (Matrix Science). The data was searched against a custom database containing the constructs of interest as well as the SwissProtein database with taxonomy restricted to *Escherichia coli*. Search parameters included trypsin digestion with up to 2 missed cleavages, peptide mass tolerance of ±10 ppm, MS/MS fragment tolerance of ±0.02 Da, fixed modification of carbamidomethyl cysteine and variable modifications of methionine oxidation and phosphorylation on serine, threonine, and tyrosine. Normal and decoy database searches were run, with the confidence level was set to 95% (p<0.05).

### Phosphotag gel assay

CK-II mediated in vitro phosphorylation of Vpu, Furin and Serinc3 was checked by Phospho-tag phosphoprotein gel stain (Genecopoeia, Rockville, MD) using the vendor-provided protocol. In brief, equal amounts of proteins that were expressed either with or without CK-II were run on an SDS-PAGE gel, which was fixed with a solution of 50% methanol, 10% acetic acid, then washed in water, before staining with the Phos-tag phosphoprotein gel stain for 90 minutes. After de-staining and washing according to the manufacturer’s instructions, the gel was imaged using a 300nm UV transilluminator.

### GST pulldown assays

Purified GST-Furin, GST-Serinc3 and MBP-µ1/MBP-µ2 proteins were used for in vitro GST-pulldown. Equimolar ratio of these proteins were mixed with GST-resin and incubated overnight at 4°C. The next day, the GST resins were extensively washed with buffer containing 20mM Tris HCL pH 7.5 and 150 mM NaCl to remove the unbound proteins. The bound proteins were eluted with 10mM glutathione reduced in 50mM Tris HCL pH 8.0. The formation of protein-protein complexes between GST-Furin or GST-Serinc3 and MBP-µ1 or MBP-µ2 were detected by SDS-PAGE using Coomassie blue stain. For “reverse” GST pulldown assays, MBP-μ2CTD-GST (0.45 mg) and MBP-SERINC3 and related mutants (∼1 mg, 5-fold molar excess) were mixed in a final volume of 100 uL. Reaction mixtures were loaded onto small, gravity flow columns containing 0.5 mL glutathione Sepharose 4B resin (GE Healthcare). The resin was extensively washed with 4 × 750 uL GST binding buffer (50 mM Tris pH 8, 100 mM NaCl, 0.1 mM TCEP). Bound protein complexes were eluted with 4× 250 uL GST elution buffer containing 10 mM reduced glutathione. Eluted proteins were analyzed by SDS-PAGE and stained with Coomassie blue.

### Size exclusion chromatography

The formation of stable protein complexes between the GST-fused CDs of Vpu, Furin and Serinc3 L10 and MBP-µ2 was assessed by size exclusion chromatography (SEC). A Superose 6 10/300GL (GE Life Sciences) small scale SEC column was used with an AKTA Pure chromatography system. The column was pre-equilibrated with buffer containing 20 mM Tris HCL pH 7.5, 150mM NaCl, 10% glycerol, 2mM DTT and 5mM EDTA. The two purified protein components (the GST-fusions and MBP-µ2) were mixed in equimolar ratio in a total volume of 100 ul, incubated at 4°C for an hour, and then injected into the SEC column. The eluting protein was detected by UV absorbance at 280nM. Molecular weight estimates for the peak fractions were derived using gel-filtration standards of various sizes. The peak fractions were analyzed by SDS-PAGE.

### Biolayer interferometry

Binding affinities between phosphorylated GST-Vpu, -Furin, and - Serinc3 with MBP-µ2 were determined by biolayer interferometry on an Octet Red instrument (ForteBio). His-tagged MBP-µ2 was immobilized on Ni-NTA capture sensors (ForteBio). The CDs of Vpu, Furin, and Serinc3, each fused to GST, were assessed as free analytes in solution (PBS at pH 7.4). For measurement of the kinetic parameters, the analytes were serially diluted to five different concentrations as shown in Figure 3. The data were analyzed using the ForteBio analysis software version 7.1, and the kinetic parameters were calculated using a global fit 1:1 model, yielding the association (k_on_), dissociation (k_off_) and affinity constants (k_d_) for each interaction.

### Phylogenetic analyses

To test whether evolution at sites 380 and 383 co-evolved with each other, we employed a Bayesian graphical model using the Spidermonkey(45) package on Datamonkey (www.datamonkey.org)(46). The relationship between the two sites was explored using a 1-parent network conditioned on a maximum liikelihood phylogeny inferred using IQ-Tree (47)under a GTR+Γ_4_ substitution model.

### HIV-1 infectivity assays

To determine the effect of SERINC3 mutations on the restriction of HIV-1 infectivity and Nef-responsiveness, 3.75×10^5^ Jurkat TAg (-*SERINC3/-SERINC5*) cells were co-transfected with either 1.15 µg pNL4-3 or pNL4-3ΔNef proviral plasmid and 100 ng plasmid encoding wild type or mutant SERINC3, using DNA-In Jurkat transfection reagent (MTI GlobalStem). Two days later, the suspension cells were pelleted by low-speed centrifugation and virions were partially purified from the culture supernates by ultracentrifugation at 20,000x*g* through a 20% sucrose cushion. The viral pellets were resuspended in Dulbecco Modified Eagle Medium (DMEM) supplemented with 10% fetal bovine serum, and infectivity was determined by infecting 2×10^4^ HeLa P4.R5 cells in a 48-well plate in duplicate with serial dilutions of the virion preparations. Two days after infection, the cells were fixed and stained for β-galactosidase activity and the number of infectious centers was measured by computer-assisted image analysis as described previously (48). The data are expressed as the number of infectious centers normalized to the concentration of HIV capsid antigen (p24), as measured by ELISA (ABL; Rockville, MD).

### Immunoblot

Cell lysates and virion pellets from infectivity experiments were analyzed by SDS– PAGE and western blotting. The viral and cell pellets were resuspended in Laemmli buffer supplemented with 50 mM TCEP. Cell samples were sonicated before SDS-PAGE and western blotting. Western blots were probed for exogenous SERINC3 expression using mouse anti-HA (HA.11, Clone 16B12, Biolegend). The blots were also probed using mouse anti-GAPDH (Genentech) as a cellular protein loading control and mouse anti-p24 (EMD Millipore) as a virion-protein loading control. Horseradish peroxidase-conjugated goat anti-Mouse IgG (BioRad) was used as a secondary antibody, and the immunoreactive bands were visualized by enhanced chemiluminescence.

## Acknowledgments

This work was supported by NIH grants to RS (CFAR supplement to P30 AI036214), JG (R37 AI081668 and R01 AI129706), YX (R01 AI102778), RW (P01 AI104722 and International AIDS Vaccine Initiative, IAVI), and JOW (K01 AI110181 and R21 AI115701). We thank Marissa Suarez for the p24 ELISAs.

## Author contributions

RS, CS, CL, JG, JOW, XJ, YX, and JG designed and/or performed the experiments. All authors interpreted the data and edited or wrote the manuscript.

